# Cell division machinery drives cell-specific gene activation during bacterial differentiation

**DOI:** 10.1101/2023.08.10.552768

**Authors:** Sylvia Chareyre, Xuesong Li, Brandon R. Anjuwon-Foster, Sarah Clifford, Anna Brogan, Yijun Su, Hari Shroff, Kumaran S. Ramamurthi

**Author notes:** The authors declare no competing interests.

## Abstract

When faced with starvation, the bacterium *Bacillus subtilis* transforms itself into a dormant cell type called a "spore". Sporulation initiates with an asymmetric division event, which requires the relocation of the core divisome components FtsA and FtsZ, after which the sigma factor σ^F^ is exclusively activated in the smaller daughter cell. Compartment specific activation of σ^F^ requires the SpoIIE phosphatase, which displays a biased localization on one side of the asymmetric division septum and associates with the structural protein DivIVA, but the mechanism by which this preferential localization is achieved is unclear. Here, we isolated a variant of DivIVA that indiscriminately activates σ^F^ in both daughter cells due to promiscuous localization of SpoIIE, which was corrected by overproduction of FtsA and FtsZ. We propose that a unique feature of the sporulation septum, defined by the cell division machinery, drives the asymmetric localization of DivIVA and SpoIIE to trigger the initiation of the sporulation program.

## INTRODUCTION

During cellular differentiation, a progenitor cell changes to assume characteristics required for a specific function. For instance, in eukaryotes, stem cells are known for their ability to differentiate into multiple cell types (2–4). Two hallmarks of this process are differential gene expression and asymmetric division, which result in two unequal daughter cells that behave dissimilarly. A relatively simple model system to study these processes is bacterial endospore formation (“sporulation”), wherein a progenitor cell differentiates into two cell types that each display a different cell fate. *Bacillus subtilis* is a gram-positive rod-shaped bacterium that normally divides by binary fission when grown in rich medium, resulting in two identical daughter cells that exhibit similar cell fates (6–8). However, when the bacterium senses the onset of starvation, it initiates the sporulation program, which results in the transformation of the cell into an ellipsoidal, dormant cell type called a “spore” that is resistant to myriad environmental assaults (9–11). Sporulation commences with an asymmetric division event near one pole of the bacterium that produces two dissimilarly sized progeny: a larger mother cell and a smaller forespore, which initially lie side-by-side, separated by the “polar septum” (Fig. 1A). The asymmetric positioning of the polar septum is achieved by redeployment of FtsZ, a bacterial tubulin homolog that is the core component of the divisome (12, 13). Redeployment requires the slight overexpression of the *ftsZ* gene (driven by a second sporulation-specific promoter upstream of the *ftsAZ* operon (14, 15), which also encodes for FtsA, an actin homolog that tethers FtsZ to the membrane (16), and a sporulation protein named SpoIIE (17, 18). Ultimately, the forespore will mature into the spore, and the mother cell will lyse after nourishing the forespore into dormancy. This transformation is driven by the sequential, compartment-specific activation of sigma factors in the forespore and mother cell (19).

**Figure 1.**
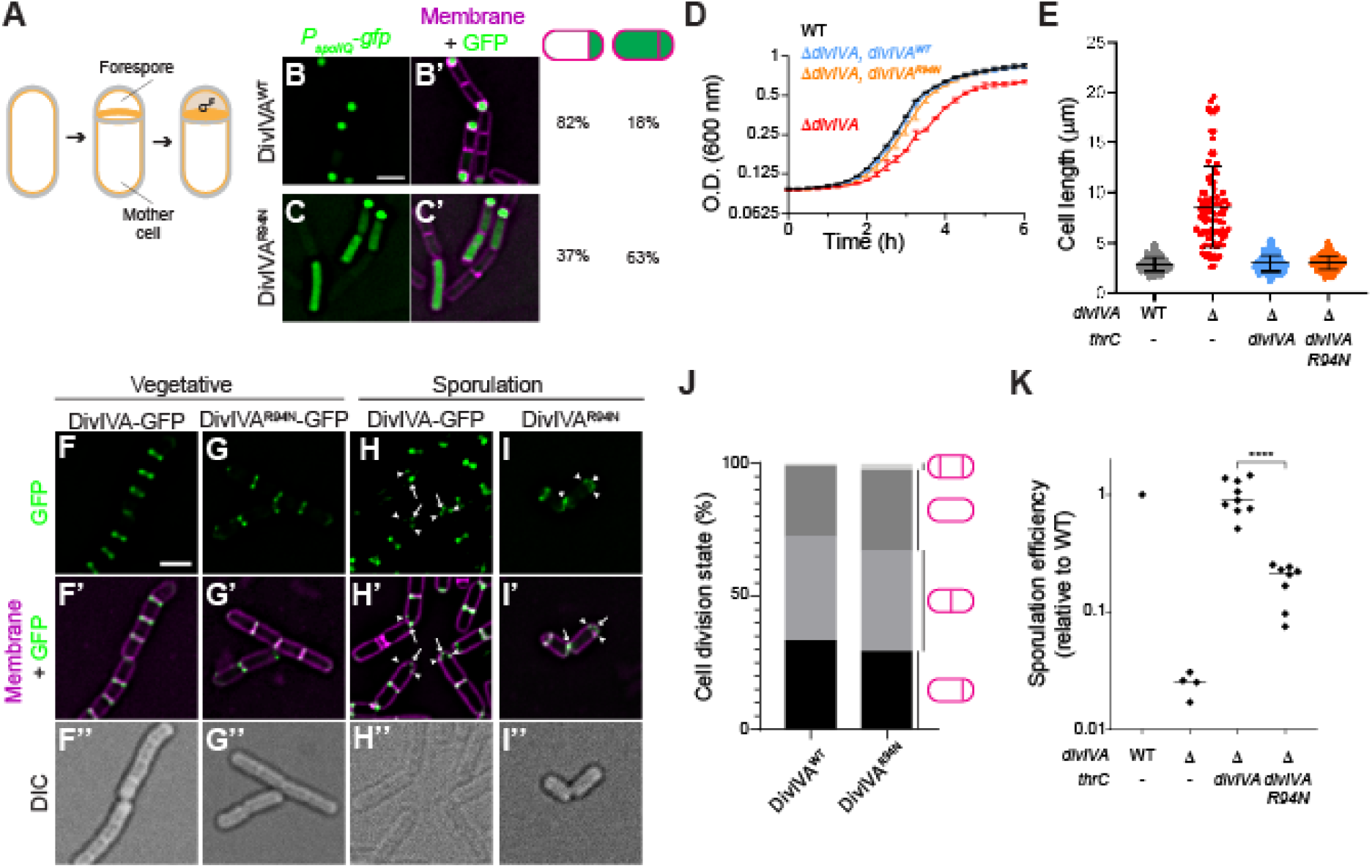
Cells expressing *divIVA^R94N^* are specifically defective for σ^F^ activation. (A) Schematic of sporulation initiation in *Bacillus subtilis*. Sporulation begins with an asymmetric division, producing a smaller “forespore” and a larger “mother cell”. The sigma factor σ^F^ is specifically activated in the forespore compartment. Membranes depicted in yellow; cell wall depicted in gray. (B-C’) Fluorescence micrographs monitoring σ^F^ activation using promoter fusions (*P*_spoIIQ_-*gfp*, a σ^F^-controlled promoter) in (B-B’) otherwise WT cells or (C-C’) cells expressing *divIVA*^R94N^ as the only copy of *divIVA* at t = 1.5 h after induction of sporulation. B-C: Fluorescence from GFP production; B’-C’: overlay, GFP and membranes visualized using FM4-64. Strains used: SJC124, SJC93. Scale bar: 1 µm. Fraction of cells (n > 200) displaying forespore-exclusive (left column) or mother cell and forespore (right column) production of GFP are shown to the right of the micrographs. (D) Growth curves of WT (black), Δ*divIVA* (red), or Δ*divIVA* strain complemented *divIVA* (blue) or *divIVA*^R94N^ (orange), as measured by optical density (O.D.) at 600 nm. Bars represent mean; errors: S.D. (n = 3 independent cultures). (E) Quantification of cell lengths WT (gray), Δ*divIVA* (red), or Δ*divIVA* strain complemented *divIVA* (blue) or *divIVA*^R94N^ (orange), measured using fluorescence microscopy with membranes visualized with FM4-64. Strains used: PY79, KR546, BRAF22, APB8. Bars represent mean; whiskers: I.Q.R. (n = 86-265 individual cells). (F-I’’) Fluorescence micrographs of otherwise WT cells producing (F-F’’, H-H’’) DivIVA-GFP, or (G-G’’, I-I’’) DivIVA^R94N^-GFP during (F-G’’) vegetative growth, or (H-I’’) 1.5 h after induction of sporulation (strains used: SC634 and SC635). (F-I) Fluorescence from GFP; (F’-I’) overlay, membranes and GFP; (F’’-I’’) differential interference contrast (DIC). Scale bar: 1 µm. Arrows indicate pole-localized DivIVA-GFP; arrowheads indicate DivIVA-GFP at the polar septum. (J) Cell morphologies of cells stained with membrane dye FM4-64 producing WT or R94N variants of DivIVA examined 1.5 h after induction of sporulation. Examined morphologies (no septa, medial septum, polar septum, or disporic) are indicated to the right and the fraction of cells exhibiting that morphology are indicated in different shades of gray. (K) Sporulation efficiencies, determined as heat resistance, of WT, Δ*divIVA*, or cells expressing *divIVA* or *divIVA*^R94N^ as the only copy of *divIVA* (strains PY79, KR546, BRAF22, APB8) and reported as relative to WT. *thrC* is an ectopic chromosomal locus from which indicated alleles of *divIVA* were expressed. Symbols are individual values from independent cultures; bars represent mean. Statistical analysis: one-way ANOVA, **** indicates P-value < 0.0001.

The first sigma factor that sets off this cascade is σ^F^, which is activated specifically in the forespore shortly after elaboration of the polar septum (20). To prevent promiscuous or premature activation, σ^F^ is held inactive by an anti-sigma factor (SpoIIAB) (21, 22). Activation of σ^F^ is achieved by dephosphorylating an anti-anti-sigma factor (SpoIIAA), which sequesters SpoIIAB and thereby liberates σ^F^ (23, 24). The phosphatase responsible for activating SpoIIAA is the FtsZ-redeploying protein SpoIIE, whose phosphatase activity has been studied in detail (25–29). A puzzling feature of this mechanism is that the three components that regulate σ^F^ are transcribed and translated in the progenitor cell before asymmetric septation, so multiple mechanisms have been proposed to explain the specific activation of σ^F^ specifically in the forespore. One model builds on the observation that SpoIIE displays a biased localization on the forespore face of the polar septum (30–32), thereby preferentially exerting its phosphatase activity on forespore-localized SpoIIAB, resulting in forespore-specific σ^F^ activation. The preferential localization of SpoIIE in the forespore is dependent on a membrane-associated structural protein called DivIVA that participates in positioning proteins during cell division and anchoring replicated chromosomes during sporulation (33), which also interacts with SpoIIE at the polar septum and likewise displays a biased localization on the forespore face of the polar septum (32). DivIVA preferentially accumulates on highly negatively curved membranes (34, 35), and as such localizes to either side of vegetative septa (5). However, the reason underlying the biased localization of DivIVA on just one side of the polar septum during sporulation, which also harbors negative membrane curvature on either side of the septum, was unclear. A second model invokes the selective degradation of SpoIIE in the mother cell, via the preferential localization of the membrane-bound protease FtsH. This model also invokes a requirement for DivIVA, but in a protective role that sequesters SpoIIE from degradation by FtsH (36).

A recent report revealed yet another level of asymmetry at the polar septum (1). Whereas DivIVA and SpoIIE localize to the forespore face of the polar septum, the core components of the cell division machinery itself (FtsA and FtsZ) reportedly display a biased localization on the mother cell face of the polar septum. This asymmetric distribution of the cell division machinery was proposed to cause the unusual thinness of the polar septum compared to vegetative septa (37, 38). In this work, we investigated how the asymmetric distribution of four different proteins on either side of a division septum is established, and if the biased localization of DivIVA at the polar septum is principally responsible for the compartment-specific activation of σ^F^. We first identified a DivIVA variant that did not display forespore-biased localization at the polar septum that triggered the promiscuous activation of σ^F^ in both the mother cell and forespore, resulted in a sporulation defect, and caused the thickness of the polar septum to resemble vegetative septa. The mis-localization of this DivIVA variant was corrected by the overproduction of FtsA and FtsZ, which restored septum thickness, forespore-biased SpoIIE localization, and proper σ^F^ activation. We propose that a unique architecture of the polar septum, resulting from the overexpression of *ftsA* and *ftsZ* at the onset of sporulation, drives the biased localization of DivIVA to establish an intrinsic asymmetry that initiates the cascade of differential transcription that drives the sporulation program.

## RESULTS

### A DivIVA variant is specifically impaired in σ^F^ activation

After elaboration of the polar septum, σ^F^ is activated specifically in the forespore, which requires a phosphatase, SpoIIE, (20, 25, 39, 40). We previously reported that selective activation of σ^F^ in the forespore is likely due to the asymmetric localization of SpoIIE phosphatase on the forespore face of the polar septum. SpoIIE directly interacts with DivIVA, which we showed, like SpoIIE, asymmetrically localizes on the forespore face of the polar septum (32). This dependence on DivIVA for SpoIIE localization was demonstrated by the controlled degradation of DivIVA at the onset of sporulation, but this approach could have also abolished other DivIVA functions that could have been concurrently occurring. To genetically separate the contribution of DivIVA in σ^F^ activation from its vegetative and chromosome anchoring roles, we mutagenized selected codons surrounding position 99 of DivIVA, which had previously been reported to specifically affect sporulation, but nonetheless had a chromosome anchoring defect (41), and screened, using fluorescence microscopy, for DivIVA variants that mis-activated σ^F^ using fluorescence microscopy. At 1.5 h after induction of sporulation, 82% of otherwise WT cells (n=272) that harbored *gfp* under control of a σ^F^-dependent promoter produced GFP exclusively in the forespore (Fig. 1B-B’). However, substituting Arg at position 94 of DivIVA with Asn (DivIVA^R94N^) resulted in production of GFP in both the mother cell and forespore in ∼60% of cells (n=227; Fig. 1C-C’). This promiscuous activation of σ^F^ resulted in a reduced sporulation efficiency: whereas cells harboring a *divIVA* deletion that was complemented with WT *divIVA* sporulated at 0.94 relative to WT, cells expressing *divIVA*^R94N^ sporulated at only 0.18 relative to WT (Fig. 1K).

We next tested if DivIVA^R94N^ was impaired in its other known functions. Unlike cells harboring a deletion of *divIVA*, cells expressing *divIVA*^R94N^ did not display an obvious exponential growth defect (Fig. 1D) and displayed a cell length that was similar to WT (Fig. 1E), suggesting that this allele was not impaired in the vegetative function of DivIVA. Consistent with this, DivIVA^R94N^-GFP primarily localized at mid-cell during vegetative growth, similar to the localization pattern of DivIVA-GFP (5) (Fig. 1F-G’’). At the onset of sporulation, DivIVA-GFP localizes to both the hemispherical cell poles and the polar septum; similarly, DivIVA^R94N^-GFP (at least as measured by diffraction-limited fluorescence microscopy) localized to the cell poles and polar septum (Fig. 1H-I’’), suggesting that there is no gross localization defect of DivIVA^R94N^ during vegetative growth or during sporulation. Degradation of DivIVA at the onset of sporulation prevents polar septation (32). To test if cells expressing *divIVA*^R94N^ displayed a defect in polar septation, we examined, using fluorescence microscopy, the morphology of cells stained with a fluorescent membrane dye 1.5 h after the induction of sporulation and quantified the fraction of cells that displayed polar or medial septa, no septa at all, or two polar septa at both poles. The results in Fig. 1J indicate that the population of cell morphologies present at this early stage of sporulation were similar between cells producing DivIVA or DivIVA^R94N^. Finally, we examined the chromosome anchoring efficiency in cells expressing *divIVA*^R94N^ by visualizing chromosomes of sporulating cells using the fluorescent dye DAPI and observing the number of forespores that were devoid of any DNA, indicating a chromosome anchoring defect. 51% of cells (n=260) harboring a deletion in *racA*, which is known to cause a chromosome anchoring defect, produced chromosome-free forespores. In contrast, only 4% of WT cells and 19% of cells expressing *divIVA*^R94N^ displayed a chromosome anchoring defect (n=191 and 311, respectively; Fig. S1A-C). To test if the σ^F^ mis-activation caused by DivIVA^R94N^ could be due to the slightly elevated chromosome anchoring defect we observed in this strain, we measured the fidelity of σ^F^ activation in cells harboring a *racA* deletion. Similar to WT, 83% of Δ*racA* cells correctly activated σ^F^ exclusively in the forespore, indicating that a chromosome anchoring defect is not responsible for promiscuous σ^F^ activation (Fig. S1D-E). Taken together, we conclude that the *divIVA*^R94N^ allele is specifically impaired in σ^F^ activation, which occurs after the previously described roles for DivIVA in vegetative growth, chromosome anchoring, and polar septation.

### DivIVA^R94N^ does not asymmetrically position SpoIIE on the forespore side of the polar septum

Since DivIVA anchors SpoIIE to the forespore face of the polar septum, we wondered if the σ^F^ activation defect could be attributed to incorrect subcellular localization of either DivIVA or SpoIIE. To distinguish between the two faces of the polar septum, which is separated by ∼80 nm, we employed dual-color 3D structured illumination microscopy (SIM) (42, 43). Additionally, to examine early events immediately after polar septation, we employed a mutant strain (Δ*spoIID*/Δ*spoIIM*) that was arrested before engulfment and thus displayed a flat septum. Further, since SpoIIE is released into the forespore and subsequently recaptured at the polar septum, we used a strain that did not produce the recapturing protein SpoIIQ to ensure that we were not detecting released, then recaptured, SpoIIE-GFP. At the onset of polar septation, WT DivIVA-GFP exhibited a biased localization to the forespore side of the invaginating septum in 72% of cells (n=67) (32) (Fig. 2A-A’’), which was evidenced by plotting the fluorescence intensity of the membrane stain relative to that of the DivIVA-GFP fluorescence to visualize the offset peak of the DivIVA-GFP intensity towards the forespore. This asymmetric positioning of DivIVA-GFP remained in cells that had completed elaboration of the polar septum (Fig. 2B-B’’). Consistent with this localization pattern, SpoIIE-GFP also displayed a forespore-biased localization in 71% (n=60) of cells during initial septal invagination (Fig. 2E-E’’), and in cells that had completed septation (Fig. 2F-F’’). In contrast, DivIVA^R94N^-GFP did not display a forespore-biased localization, with 54% of cells displaying DivIVA^R94N^-GFP fluorescence that overlapped with the membrane fluorescence (Fig. 2C-D’’). Concomitantly, in 66% of cells harboring *divIVA*^R94N^ (n=54), SpoIIE-GFP also did not asymmetrically localize to the forespore face of the polar septum (Fig. 2G-H’’). The results suggest that the inability of DivIVA^R94N^ to display a biased localization on the forespore face of the polar septum leads to improper positioning of SpoIIE, which leads to the promiscuous activation of σ^F^ in both compartments. We wondered if this mis-localization of SpoIIE in the presence of DivIVA^R94N^, was due to a reduced interaction between both proteins. Since DivIVA and SpoIIE interact weakly in vitro (32, 36), we measured the interaction between these proteins using a two-hybrid assay in which SpoIIE and DivIVA were fused to the T18 and T25 subunits of the adenylate cyclase, respectively, and heterologously produced in *E. coli*. SpoIIE-T25 and DivIVA^WT^-T18 interacted in this assay, as evidenced by increased β-galactosidase activity (Fig 2I, lanes 1-2). By comparison, the interaction of SpoIIE-T25 with DivIVA^R94N^-T18 showed a similar β-galactosidase activity (Fig 2I, lane 3), suggesting that the R94N substitution did not appreciably alter the interaction between DivIVA and SpoIIE. Similarly, the R94N substitution also did not appreciably affect DivIVA self-interaction, since we observed similar β-galactosidase activities between DivIVA^WT^-T25 and DivIVA^WT^-T18, and DivIVA^WT^-T25 and DivIVA^R94N^-T18 (Fig 2I, lanes 4-6). The results suggest that the R94N substitution affects an intrinsic ability of DivIVA to asymmetrically localize to the forespore face of the polar septum, in a manner that does not depend on the interaction between DivIVA and SpoIIE.

**Figure 2.**
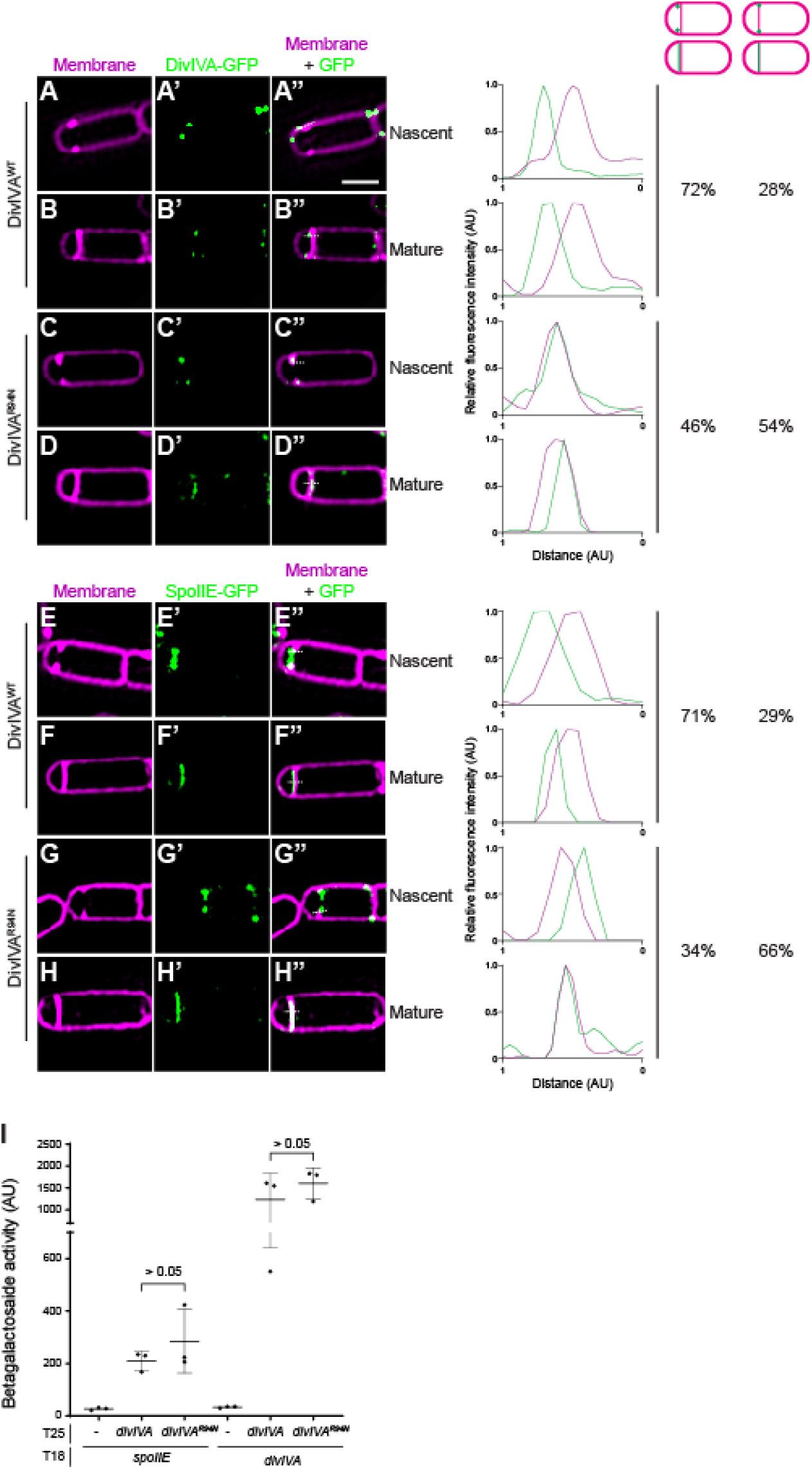
DivIVA and SpoIIE subcellular localization is impaired in DivIVA^R94N^, but interaction between DivIVA and SpoIIE is not. (A-H’’) Subcellular localization of (A-D’’) DivIVA-GFP or (E-H’’) SpoIIE-GFP monitored using dual-color 3D structured illumination microscopy in mutant cells that are blocked before the engulfment stage of sporulation (Δ*spoIIM* Δ*spoIID*), imaged 2 h after induction of sporulation, displaying (A-A’’, C-C’’, E-E’’, G-G’’) nascent or (B-B’’, D-D’’, F-F’’, H-H’’) mature septa, and producing (A-B’’, E-F’’) DivIVA^WT^, or (C-D’’, G-H’’) DivIVA^R94N^. A-H: membranes visualized using FM-64; A’-H’: fluorescence from GFP; A’’-H’’: overlay, membrane and GFP. Strains: SC634, SC635, SC656, SC657. Scale bar: 1 µm. Line scan analyses of normalized fluorescence intensity from GFP (green trace) or membrane stain (magenta trace) along the axis of the dashed line (indicated in A’’-H’’) in both channels at the selected polar septa are shown at the right, and percentage of cells that exhibit forespore-biased GFP localization (left column) or septum co-localized GFP localization (right column) are indicated. (I) β-galactosidase activity in a bacterial 2-hybrid assay between SpoIIE-T18 or DivIVA-T18 with DivIVA-T25 or DivIVA^R94N^-T25. Data points represent individual values of three biological replicates; bars represent mean; errors: S.D.

### FtsA and FtsZ influence DivIVA and SpoIIE placement

The localization pattern for DivIVA and SpoIIE is the opposite of what was recently reported for the central divisome components FtsA and FtsZ at the polar septum (44). Whereas DivIVA and SpoIIE display a forespore-biased localization, FtsA and FtsZ preferentially localize to the mother cell face of the polar septum (Fig. 3A). Moreover, the redeployment of FtsA and FtsZ to polar cell division sites itself requires SpoIIE, which directly interacts with FtsZ polymers at the onset of sporulation (45–47). We therefore wondered if FtsA and FtsZ could reciprocally impact DivIVA^R94N^ and SpoIIE placement at the polar septum. Since the *ftsAZ* operon is upregulated at the onset of sporulation (14), we tested if the slight additional overproduction of FtsA and FtsZ by an extra copy of *ftsAZ* engineered at an ectopic chromosomal site could correct the sporulation defect caused by DivIVA^R94N^. Whereas cells producing DivIVA^R94N^ sporulated with an efficiency of 0.18 relative to WT, production of FtsA and FtsZ from the second locus increased sporulation efficiency in this strain to 0.70 relative to WT (Fig. 3B, lanes 4-5). The overproduction of FtsA and FtsZ also corrected the σ^F^ activation defect caused by DivIVA^R94N^: in cells expressing *divIVA*^R94N^, only 40% of cells displayed forespore-specific activation of σ^F^, but slight overproduction of FtsAZ in this strain restored proper σ^F^ activation to 78%, similar to WT levels (Fig. 3C). However, overproduction of FtsA or FtsZ alone did not correct the sporulation defect caused by DivIVA^R94N^ (Fig. 3B, lanes 6-7), or restore forespore-specific activation of σ^F^ (Fig. 3C), indicating a combined requirement for FtsA and FtsZ overproduction to suppress the DivIVA^R94N^ defects.

**Figure 3.**
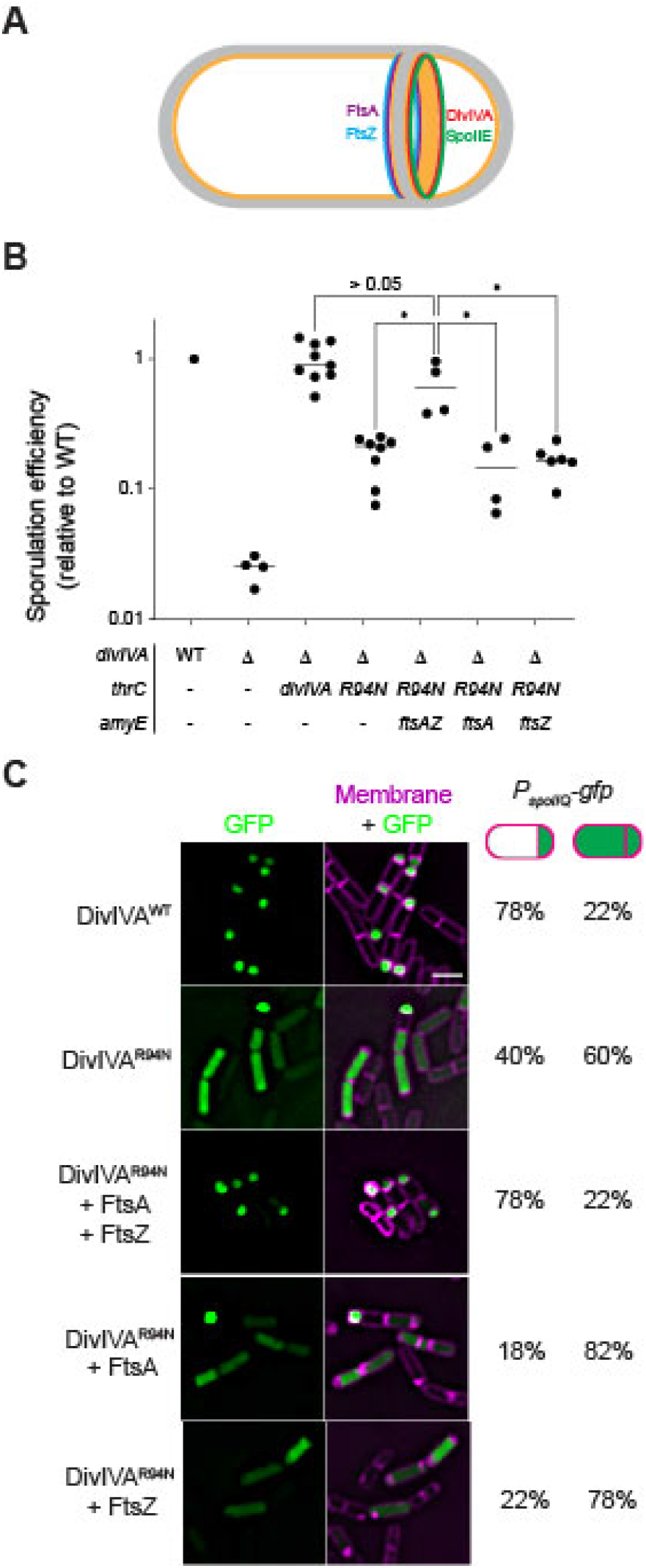
Overproduction of FtsA and FtsZ corrects the sporulation defect of DivIVA^R94N^. (A) Schematic of mother-cell biased localization of FtsA and FtsZ (purple and blue, respectively) (1), and forespore-biased localization of DivIVA and SpoIIE (red and green, respectively) at the polar septum at the onset of sporulation. (B) Sporulation efficiencies, determined as heat resistance, of WT, Δ*diviVA*, or cells expressing *divIVA* or *divIVA*^R94N^ while overexpressing *ftsAZ*, *ftsA*, or *ftsZ* from the ectopic chromosomal locus *amyE* (strains PY79, KR546, BRAF22, APB8, SJC112, SC527, SC544) and reported as relative to WT. Symbols are individual values from independent cultures; bars represent mean. Statistical analysis: one-way ANOVA, * indicates P-value < 0.05. (C) Fluorescence micrographs monitoring σ^F^ activation using promoter fusions (*P*_spoIIQ_, a σ^F^-controlled promoter) in presence of indicated DivIVA variant, with overexpression of *ftsAZ*, or *ftsA* and *ftsZ* alone. Left panels: fluorescence from GFP; right panels: overlay, GFP and membranes visualized using FM4-64. Strains: SJC124, SJC93, SJC125, SC529, SC546. Scale bar: 2 µm. Fraction of cells (n > 40) displaying forespore-exclusive (left column) or mother cell and forespore (right column) production of GFP are shown to the right of the micrographs.

We next tested if the slight overproduction of FtsAZ could correct the mis-localization of DivIVA^R94N^ and SpoIIE at the polar septum using dual-color 3D SIM. DivIVA^R94N^-GFP displayed forespore-biased localization in only 46% of cells (n=126; Fig. 4B-B’’), compared to WT DivIVA, which preferentially localized to the forespore side of the polar septum in 72% of cells (n=44; Fig. 4A-A’’). However, overproduction of FtsA and FtsZ restored DivIVA^R94N^-GFP to the forespore face of the polar septum to near WT levels (68%; Fig. 4C-C’’). We next observed the localization of SpoIIE-GFP. In the presence of WT DivIVA, 71% (n=60) of cells displayed SpoIIE-GFP on the forespore face of the polar septum (Fig. 4D-D’’); in the presence of DivIVA^R94N^, only 33% (n=56) displayed preferential localization of SpoIIE-GFP on the forespore (Fig. 4E-E’’). Overproduction of FtsA and FtsZ, though, restored the forespore-biased localization of SpoIIE-GFP in 64% (n=65) of cells, despite these cells harboring DivIVA^R94N^ (Fig. 4F-F’’). The data are consistent with a model in which SpoIIE initially triggers the redeployment of FtsA and FtsZ to polar positions (14), but reciprocally, FtsA and FtsZ drive the asymmetric distribution of SpoIIE on one face of the polar septum.

**Figure 4.**
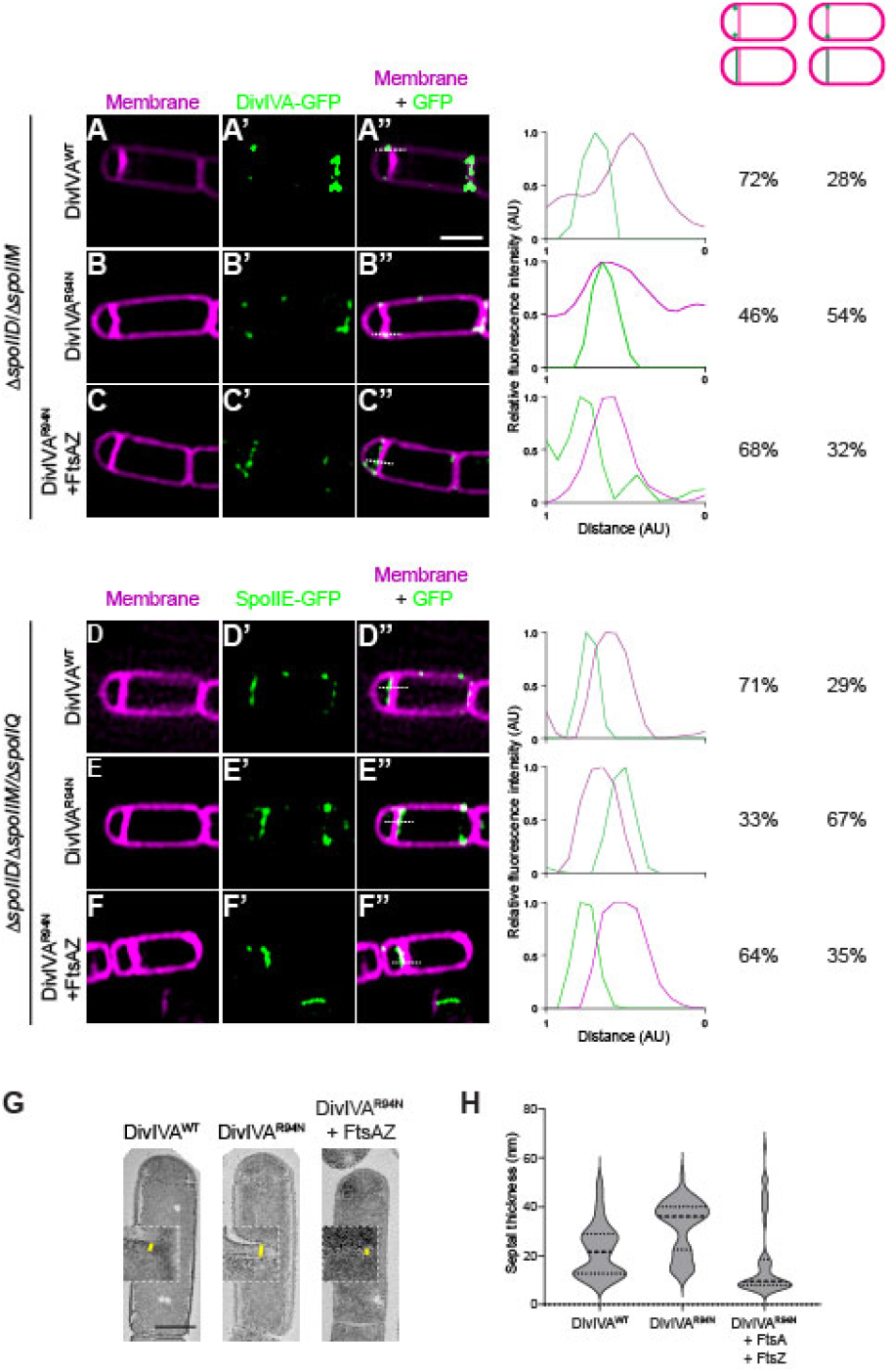
A second copy of *ftsAZ* restores mother cell-biased localization of DivIVA^R94N^ and SpoIIE as well as septum thinness. Subcellular localization of (A-C’’) indicated DivIVA-GFP variant or (D-F’’) SpoIIE-GFP monitored using dual-color 3D structured illumination microscopy in mutant cells that are blocked before the engulfment stage of sporulation (Δ*spoIIM* Δ*spoIID*), in the presence of (A-B’’, D-E’’) one copy of *ftsAZ* at the native locus, or (C-C’’, F-F’’) in cells harboring a second copy of *ftsAZ* at an ectopic locus, imaged 2 h after induction of sporulation. Cells in (D-F’’) harbor an additional deletion of *spoIIQ*, which prevents the recapture of SpoIIE at the polar septum. A-F: membranes visualized using FM-64; A’-F’: fluorescence from GFP; A’’-F’’: overlay, membrane and GFP. Strains: SC634, SC635, SC688, SC656, SC657, SC671. Scale bar: 1 µm. Line scan analyses of normalized fluorescence intensity from GFP (green trace) or membrane stain (pink trace) along the axis of the dashed line (indicated in A’’-F’’) in both channels at the selected polar septa are shown at the right, and percentage of cells that exhibit forespore-biased GFP localization (left column) or septum co-localized GFP localization (right column) are indicated. (G) Septum thickness of strains harboring DivIVA^WT^, DivIVA^R94N^, and DivIVA^R94N^ harboring an extra copy of *ftsAZ* (strains BRAF22, APB8 and SJC112) observed by transmission electron microscopy. Scale bar: 500 nm. (H) Measurements of septal thickness of at least 3 cells per strain. Violin plot represents 10 measurements per cell taken along the length of the septum.

Khanna *et al*. reported that the asymmetric positioning of FtsA and FtsZ is responsible for the unusual thinness of the polar septum as compared to vegetative septa (1). Since FtsA and FtsZ influence DivIVA and SpoIIE positioning, we wondered if DivIVA^R94N^ impacts the thickness of the polar septum. We therefore examined the effect of DivIVA^R94N^ on polar septum thickness using transmission electron microscopy. Similar to what was reported previously, we observed that the thickness of the polar septum in cells producing WT DivIVA was 20 nm (IQR = 45.0 nm) (Fig. 4H). In contrast, cells producing DivIVA^R94N^ displayed thicker polar septa of 36 nm (IQR = 40.5 nm) which, although is thinner than ∼80 nm vegetative septa, was nonetheless thicker than a WT polar septum. The overproduction of FtsA and FtsZ, though, largely corrected the thick polar septum defect caused by DivIVA^R94N^ and decreased polar septum thickness to 10 nm (IQR = 58.5 nm). In sum, the results indicate that FtsA and FtsZ influence the correct localization of the DivIVA/SpoIIE complex at the polar septum, likely by changing the architecture of the polar septum (which is manifested at one level by thinness of this septum). Additionally, the observation that DivIVA^R94N^ influences polar septum thickness suggests that the DivIVA/SpoIIE complex reciprocally influences the function of FtsA and FtsZ at the polar septum.

## DISCUSSION

DivIVA is a scaffolding protein that performs multiple functions in *B. subtilis* at different points of the cell cycle (33). In this study, we examined an understudied role of DivIVA near the onset of sporulation: the asymmetric tethering of the phosphatase SpoIIE on the forespore face of the polar septum to achieve compartment-specific activation of σ^F^ in the forespore. Previous studies reported the asymmetric localization of DivIVA and SpoIIE at the polar septum immediately after membrane invagination commenced (32), followed by the preferential release of SpoIIE into the forespore (33), where it is protected from proteolysis by the mother cell localized protease FtsH (36). However, it was not clear if the asymmetric distribution of DivIVA and SpoIIE was simply coincidental with compartment-specific gene expression in the forespore, and if degradation by FtsH produced the asymmetric distribution of DivIVA and SpoIIE at the polar septum. Based on our genetic and cytological data, we propose that components of the divisome, combined with some unique feature of the sporulation polar septum that is absent in vegetative, medial septa, drive the asymmetric localization of the DivIVA/SpoIIE complex to the forespore face of the polar septum and that this initial asymmetric localization is the primary driver of compartmentalized gene expression in the forespore.

This model is consistent with two principal observations. First, a DivIVA variant, DivIVA^R94N^, promiscuously localized to both sides of the polar septum, and along with it tethered SpoIIE to either side of the polar septum, which resulted in the activation of σ^F^ in both mother cell and forespore compartments. Thus, despite the presence of FtsH in the mother cell compartment, the initial mis-localization of SpoIIE to the mother cell face of the polar septum was sufficient to activate σ^F^ in the mother cell and cause a reduction in sporulation efficiency. Second, correction of this promiscuous mis-localization of the DivIVA^R94N^/SpoIIE complex was achieved by overproducing two core components of the cell division machinery: FtsZ and FtsA. This suggested that the process of asymmetric cell division itself was linked to the biased localization of DivIVA and SpoIIE. Moreover, the previously described role of SpoIIE in driving the polar localization of FtsZ to achieve asymmetric cell division (17), indicates a reciprocal dependence of DivIVA/SpoIIE and FtsA/FtsZ for proper localization of each complex.

DivIVA preferentially binds to highly negatively curved membranes (48). Consistent with this notion, during vegetative growth, DivIVA localizes to both sides of the division septum and forms two static rings that abut the septum on either side (5). During sporulation, DivIVA initially localizes similarly to both sides of the nascent polar septum (32), but strangely this double ring pattern is not stably maintained in mature polar septa, despite the presence of negative curvature on either side of this septum (32). What, then, distinguishes the polar septum from a vegetative septum? Other than subcellular placement, a major difference is that the polar septum is much thinner than the vegetative septum, suggesting a fundamentally different architecture (38). Recently, Khanna et al. reported that this reduction in septum thickness is driven by the asymmetric localization of FtsA and FtsZ on the mother cell face of the polar septum, which in turn requires SpoIIE (1). We therefore propose an integrated model (Fig. 5) in which SpoIIE initially triggers the redeployment of FtsA/FtsZ to polar positions in the cell and places the divisome on the mother cell face of the invaginating septum, which results in a thin septum (Fig. 5 top right cell) (1). At the onset of polar cell division, membrane invagination recruits DivIVA to the elaborating polar septum, where DivIVA binds to SpoIIE. However, the presence of the divisome on the mother cell face excludes DivIVA from the mother cell face of the polar septum, perhaps due to a unique architecture of a thin septum that is produced at the polar position and/or due to crowding by FtsA and FtsZ that prevents DivIVA from localizing on the mother cell side. This exclusion of DivIVA is somehow overcome by the DivIVA^R94N^ variant (which produces a thick polar septum, likely due to the mis-localization of the divisome on either side of the polar septum). Presumably, the overexpression of *ftsAZ* corrects the DivIVA^R94N^ defect by re-establishing a thin polar septum and restores the asymmetric localization of FtsA/FtsZ and DivIVA/SpoIIE on either side of the polar septum. Finally, we propose that any remaining SpoIIE on the mother cell face of the polar septum is degraded by FtsH (36) to ensure the exclusive localization of SpoIIE in the forespore to activate σ^F^ in that compartment. Thus, the elaboration of an unusual division septum, driven by an interplay between divisome components and a cell fate determinant, establishes intrinsic asymmetry to trigger cellular differentiation.

**Figure 5.**
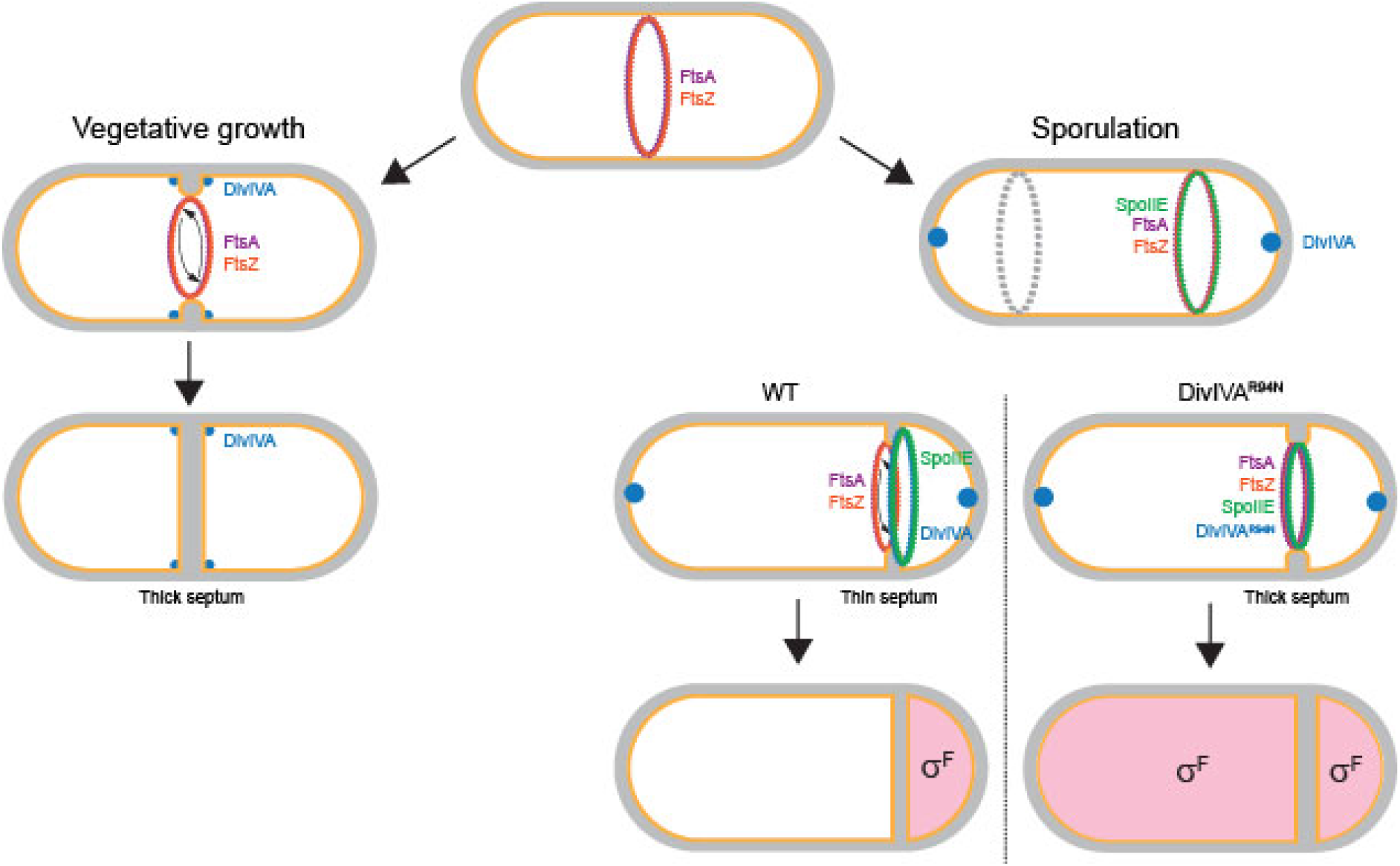
Model for compartment-specific activation of σ^F^. Schematic of *Bacillus subtilis* divisome placement in during (left) vegetative growth and (right) sporulation. FtsA (purple) and FtsZ (orange) rings initially assemble at mid-cell. During vegetative growth, the divisome constricts, creating highly negatively curved membrane on either side of the division septum, which recruits DivIVA (blue), which forms double rings that flank the septum (depicted as dots). Once constriction completes, DivIVA remains abutted to either side of the mature septum (5). During sporulation, DivIVA localizes to the extreme poles and SpoIIE (green) associates with FtsZ at mid-cell and drives the redeployment of FtsA and FtsZ to the poles of the bacterium, whereupon only one FtsZ ring actively constricts to form the polar septum and the other (gray) remains inactive. In a WT cell, FtsA and FtsZ mediate septation, but localize to the mother cell face of the septum, resulting in a unique septal architecture that is manifested, in part, as a thin septum. DivIVA and SpoIIE display a forespore-biased localization at this septum, resulting in preferential activation of σ^F^ in the forespore compartment (depicted in pink). Cells producing DivIVA^R94N^ disrupt the biased localization of FtsA and FtsZ, resulting in the elaboration of a more vegetative-like septum (that is in part manifested as a thicker septum), and in turn results in the unbiased localization of DivIVA ^R94N^ and SpoIIE at the polar septum, resulting in the promiscuous activation of σ^F^ in both compartments.

## ACKNOWLEDGEMENTS

We thank S. Gottesman, G. Storz, S. Wickner, A. Khare, M. Maurizi, N. Bradshaw, and I. Barák for discussions; P. Eswara, and members of the K.S.R. laboratory for comments on the manuscript; and C. Burks and F. Soheilian of the Electron Microscopy Laboratory of CCR for TEM sample preparation and imaging. This work was funded by the Intramural Research Program of the National Institutes of Health, Advanced Imaging and Microscopy Resource (J.C. and H.S.); the National Institute of Biomedical Imaging and Bioengineering (H.S.); and the National Cancer Institute, Center for Cancer Research (K.S.R.). This work was supported by the Howard Hughes Medical Institute (HHMI). This article is subject to HHMI’s Open Access to Publications policy. HHMI laboratory heads have previously granted a non-exclusive CC BY 4.0 license to the public and a sub-licensable license to HHMI in their research articles. Pursuant to those licenses, the author-accepted manuscript of this article can be made freely available under a CC BY 4.0 license immediately upon publication.

## MATERIALS AND METHODS

### Strain construction and general methods

All *Bacillus subtilis* strains are derivatives of *Bacillus subtilis* PY79 (49); genotypes are listed in Table S1. Genes of interest were amplified using PCR to include their native promoter and cloned into integration vectors pDG1662 (for insertion into the *amyE* locus) or pDG1731 (for insertion into the *thrC* locus) (50, 51). The *divIVA^R94N^* allele was generated using the QuikChange site-directed mutagenesis kit (Agilent). For *B. subtilis* growth curves, one colony on each strain were first inoculated in 2 ml Difco Sporulation Medium (DSM; KD Medical). 150 µl of this suspension was then placed in individual wells of a 96-well plate. Cells were grown at 37 °C for 6 h, with shaking, using a microtiter plate reader (Tecan) and the optical density at 600 nm was measured every 15 min. All *Escherichia coli* strains and plasmids used in this publication are listed in Table S1.

### Sporulation efficiency assay

Strains were grown in DSM for 24 h at 37 °C at 250 rpm and were then subjected to 80 °C for 20 min in a water bath to kill nonsporulating cells and defective spores. Serial dilutions in phosphate buffered saline were plated on LB agar, incubated at 37 °C overnight. Sporulation efficiency was determined by enumerating colony forming units (cfu) per ml and reported relative to cfu obtained for a culture of WT (strain PY79) grown in parallel.

### Epifluorescence microscopy

Cells were induced to sporulate using the resuspension method in SM medium (52). At different time points, 150 µl of cell cultures were harvested and resuspended in 10 µl PBS containing 5 µg ml^-1^ FM4-64 membrane dye and/or 2 µg ml^-1^ DAPI to visualize DNA, as needed. Finally, 3 µl of cell suspension were placed on a glass bottom culture dish (MatTek) and covered with 1 % agar pad made with SM medium. Cells were viewed using a DeltaVision core microscope system equipped with an environmental chamber at 22 °C. Cells images were captured with a Photometrics cool snap HQ2 camera. Eights planes were acquired every 0.2 µm and the data were submitted to deconvolution using SoftWorx software (53).

### Dual-color 3D structured illumination microscopy

Super-resolution imaging was conducted on a custom-built 4-beam SIM system equipped with two lasers (488 nm and 561 nm, Coherent, Sapphire 488 LP-300 mW and Sapphire 561 LP-200 mW), a phase-only nematic spatial light modulator (SLM; Meadowlark Optics, MSP1920-400-800-HSP8), a water objective lens (Nikon, CFI SR Plan Apo ×60/1.27 NA) and a piezo z-stage (Applied Scientific Instrumentation, PZ-2150, 150-µm axial travel) (54). In this work, only the 3D SIM acquisition mode was used. High-precision #1.5 coverslips (Thorlabs, CG15XH) were cleaned by immersion in 75% ethanol overnight and air dried before use. To ensure effective adherence of bacteria to the coverslips, a droplet of 10 µL poly-L-lysine solution (Sigma-Aldrich, P8920) was applied to the center of each coverslip in a biosafety cabinet. The solution was air dried at room temperature, after which the coverslips were rinsed with pure ethanol and left to air dry until ready for use. Orange FluoSpheres (Invitrogen, F8800, 0.1-µm diameter) were used as the fiducials for correcting local chromatic aberrations in dual-color imaging. Orange beads were dissolved in pure methanol at 1:50,000 dilution, and 2 µl of the solution was applied to the center of the poly-L-lysine-coated coverslips immediately before use. Beaded coverslips were mounted in a magnetic imaging chamber (Warner Instruments, QR-40LP) for imaging. After staining membranes with FM4-64, cells were washed three times with 1× PBS, each wash centrifuging the solution at 3000 rpm for 3 min. Cells were concentrated in 100 µl of 1× PBS stock solution and a 2 µl droplet of the stock was applied to the center of a beaded coverslip. Cells were allowed to settle for 1 min and the sample was rinsed once by 1× PBS before imaging. To minimize photobleaching, the FM4-64 dye was excited at 561 nm, followed by excitation of the green fluorescent protein at 488 nm. To estimate and correct chromatic aberrations in the system, the apparent position of the orange beads in both color channels was recorded and used to register the images. Image registration was conducted in ImageJ so that the orange beads colocalized in both lateral and axial views for each segmented region of interest.

Images were collected over 2 µm (ensuring we imaged the entire thickness of each bacterium) with an axial step size of 0.125 µm. We used an exposure time of 50 ms per phase, resulting in 27 s imaging time for dual-color volume acquisition. The approximate intensity at the sample plane was 80 W/cm^2^ and 60 W/cm^2^ for 488 nm and 561 nm, respectively. Further details on 3D SIM reconstruction algorithms and associated software have been described (54). Following reconstruction, all SIM images had a lateral pixel size of 40.9 nm. Microscopy images represent single plane and the asymmetric positioning of DivIVA and SpoIIE were highlighted on a graph using line scan tool using Fiji software. Arbitrary axes were chosen in the area where both the GFP and the membrane signals were the strongest.

### Bacterial two-hybrid assay

Plasmids harboring *spoIIE* or *divIVA* are derived from pKNT25 and pUT18 and were co-transformed in *E. coli* BTH101 strain (55). Transformants were plated on LB agar containing 100 μg mL^-1^ ampicillin, 50 μg mL^-1^ kanamycin and 1% glucose, and incubated at 30 ⁰C for 96 h. For each co-transformation, 10 colonies were pooled together and grown overnight at 30 ⁰C with shaking in LB medium containing ampicillin, kanamycin, and 1 mM IPTG. Cells were then diluted 1:10 into fresh LB medium in a 96 well-plate and OD_600_ was measured using a plate reader (Tecan). 100 µl of the diluted culture were placed in a new 96 well plate and lysed by addition of 10 μl of Lysis Buffer (1 mg mL^-1^ lysozyme in 1X BugBuster buffer (Sigma)) at room temperature for 15 min. 100 µl of Buffer Z (62 mM Na_2_HPO_4_, 45 mM NaH_2_PO_4_, 10 mM KCl, 1 mM MgSO_4_, 50 mM β-mercaptoethanol) was added and the beta-galactosidase reaction was started by adding 2 mM (final concentration) ortho-Nitrophenyl-β-galactoside (ONPG). Hydrolysis of ONPG was monitored by measuring A_420_ every 5 seconds for 30 min using a microplate reader (Tecan). Β-galactosidase activity was calculated by measuring the *V*_max_ of the A_420_ appearance divided by the OD_600_. Values were then multiplied by 100000, a coefficient that was chosen empirically to approximate Miller units.

### Transmission electron microscopy

Strains were induced to sporulate using the resuspension method. At t = 1.5 h cells were harvested by centrifugation and fixed in 4% formaldehyde, 2% glutaraldehyde in 0.1 M cacodylate buffer, post fixed using a 1% osmium tetroxide solution, then dehydrated sequentially in 35%, 50%, 75%, 95% and 100% ethanol followed by 100% propylene oxide. Cells were infiltrated in an equal volume of 100% propylene oxide and epoxy resin overnight and embedded in pure resin the following day. The epoxy resin was cured at 55 °C for 48 h. The cured block was thin-sectioned and stained in uranyl acetate and lead citrate. The sample was imaged with a Hitachi H7600 TEM equipped with a CCD camera (56). Septum thickness measurements were taken using Fiji software. The line tool was used to evaluate the distance over the septum, and each value was plotted in a violin graph.

## SUPPLEMENTAL MATERIAL

**Figure S1.**
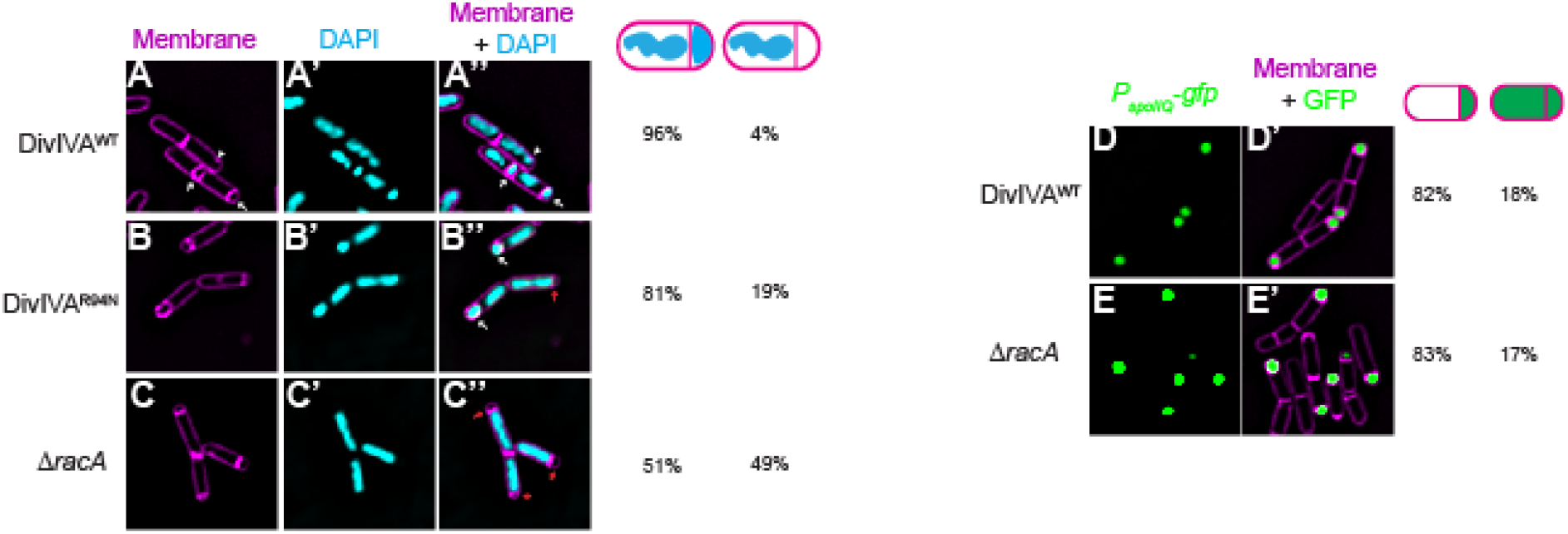
DivIVA^R94N^ does not cause a defect in chromosome anchoring at the onset of sporulation. (A) Fluorescence micrographs of sporulating cells of *B. subtilis* producing (A-A’’) WT DivIVA, (B-B’’) DivIVA^R94N^, or (C-C’’) harboring a deletion of *racA* imaged 1.5 h after the onset of sporulation. (A-C) Membranes visualized using FM4-64; (A’-C’) chromosomes visualized using DAPI; (A’’-C’’) overlay, membranes and chromosome. (E-E’) Fluorescence micrographs monitoring σ^F^ activation using promoter fusions (*P*_spoIIQ_-*gfp*, a σ^F^-controlled promoter) in (D-D’) otherwise WT cells or (E-E’) cells harboring deletion of *racA* at t = 1.5 h after induction of sporulation. D-E: Fluorescence from GFP production; D’-E’: overlay, GFP and membranes visualized using FM4-64.

**Table S1.** *Bacillus subtilis, Escherichia coli* strains and plasmids used in this study.

| Strains |  |  |
| --- | --- | --- |
| Name | Genotype | Source |
| PY79 | Prototrophic derivative of *B. subtilis* 168 | (1) |
| KR546 | $\Delta divIVA::erm$ | (2) |
| BRAF22 | $\Delta divIVA::erm thrC::divIVA spec$ | This study |
| APB8 | $\Delta divIVA::erm thrC::divIVA^{R94N} spec$ | This study |
| SJC124 | $\Delta divIVA::cm thrC::divIVA spec sacA::P_{spolIQ}-GFP kan$ | This study |
| SJC93 | $\Delta divIVA::cm thrC::divIVA^{R94N} spec sacA::P_{spolIQ}-GFP kan$ | This study |
| SC634 | $\Delta divIVA::erm \Delta spolID \Delta spolIM thrC::divIVA spec amyE::divIVA GFP cat$ | This study |
| SC635 | $\Delta divIVA::erm \Delta spolID \Delta spolIM thrC::divIVA^{R94N} spec amyE::divIVA^{R94N}-GFP cat$ | This study |
| SC656 | $\Delta divIVA::erm \Delta spolID \Delta spolIM \Delta spolIQ thrC::divIVA spec spolIE-GFP kan$ | This study |
| SC657 | $\Delta divIVA::erm \Delta spolID \Delta spolIM \Delta spolIQ thrC::divIVA^{R94N} spec spolIE-GFP kan$ | This study |
| SJC112 | $\Delta divIVA::erm thrC::divIVA^{R94N} spec amyE::P_{ftsA}-ftsAZ cat$ | This study |
| SJC125 | $\Delta divIVA::erm thrC::divIVA^{R94N} spec sacA::P_{spolIQ}-GFP kan amyE::P_{ftsA}-ftsAZ cat$ | This study |
| SC527 | $\Delta divIVA::erm thrC::divIVA^{R94N} spec amyE::P_{ftsA}-ftsA cat$ | This study |
| SC529 | $\Delta divIVA::erm thrC::divIVA^{R94N} spec sacA::P_{spolIQ}-GFP kan amyE::P_{ftsA}-ftsA cat$ | This study |
| SC544 | $\Delta divIVA::erm thrC::divIVA^{R94N} spec amyE::P_{ftsA}-ftsZ cat$ | This study |
| SC546 | $\Delta divIVA::erm thrC::divIVA^{R94N} spec sacA::P_{spolIQ}-GFP kan amyE::P_{ftsA}-ftsZ cat$ | This study |
| SC688 | $\Delta divIVA::erm \Delta spolID \Delta spolIM thrC::divIVA^{R94N} spec amyE::divIVA^{R94N}-GFP cat sacA::ftsAZ cat::tet$ | This study |
| SC671 | $\Delta divIVA::erm \Delta spolID \Delta spolIM thrC::divIVA^{R94N} spec spolIE-GFP kan sacA::ftsAZ cat::tet$ | This study |
| BTH101 | *E. coli* F', *cya*-99, *araD*139, *galE*15, *galK*16, *rpsL*1 (*StrR*), *hsdR*2, *mcrA*1, *mcrB*1, *relA*1 Euromedex (ref. EUK001) | (3) |
| Plasmids |  |  |
| Name | Description | source |
| pKNT25 | Derived from pSU40 Plac-MCS(HindIII-SphI-PstI-XbaI-BamHI-SmaI-KpnI-SacI-EcoRI)-T25 | (3) |
| pUT18 | Derived from pUC19. Plac-MCS(HindIII-SphI-PstI-Sall-XbaI-BamHI-SmaI-KpnI-SacI-EcoRI)-T18 | (3) |
| pSC0209 | SpolIE-T18 derived from pUT18 | This study |
| pSC0207 | DivIVA<sup>WT</sup>-T18 derived from pUT18 | This study |
| pSC0210 | DivIVA<sup>WT</sup>-T25 derived from pKNT25 | This study |
| pSC0211 | DivIVA<sup>R94N</sup>-T25 derived from pKNT25 | This study |

- 540 1. Youngman P, Perkins JB, Losick R. Construction of a cloning site near one end of Tn917 into which foreign DNA may be inserted without affecting transposition in *Bacillus subtilis* or expression of the transposon-borne erm gene. *Plasmid*. 1984;12: 1-9. - 543 2. Eswaramoorthy P., Erb LM, Gergory AJ, Silverman J, Pogliano K, Pogliano J and Ramamurthi KS, Cellular architecture mediates DiviVA ultrastructure and regulates min activity in *bacillus subtilis*. *mBio* 2011 - 545 3. Battesti and Bouveret, the bacterial two-hybrid system based on adenylate cyclase reconstitution in Escherichia coli, *Methods*,

## Notes

### Competing Interest Statement

The authors have declared no competing interest.

